# Short residence times of DNA-bound transcription factors can reduce gene expression noise and increase the transmission of information in a gene regulation system

**DOI:** 10.1101/776955

**Authors:** Eugenio Azpeitia, Andreas Wagner

## Abstract

Gene expression noise is not just ubiquitous but also variable, and we still do not understand some of the most elementary factors that affect it. Among them is the residence time of a transcription factor (TF) on DNA, the mean time that a DNA-bound TF remains bound. Here, we use a stochastic model of transcriptional regulation to study how this residence time affects gene expression. We find that the effect of residence time on gene expression depends on the level of induction of the gene. At high levels of induction, residence time has no effect on gene expression. However, as the level of induction decreases, short residence times reduce gene expression noise. The reason is that fast on-off TF binding dynamics prevent long periods where proteins are predominantly synthesized or degraded, which can cause excessive fluctuations in gene expression. As a consequence, short residence times can help a gene regulation system acquire information about the cellular environment it operates in. Our predictions are consistent with the observation that experimentally measured residence times are usually modest and lie between seconds to minutes.

## Introduction

All gene expression is noisy. It produces mRNA and protein molecules whose numbers fluctuate randomly. Such noise is caused by stochastic molecular interactions, which include interactions between transcription factors and DNA, and by the stochastic synthesis and degradation of molecules (1,2). Gene expression noise affects multiple biological processes. For example, it can promote phenotypic diversity, influence the coordination of gene expression, trigger cell differentiation, and facilitate evolutionary transitions (1–7). Furthermore, noise can also reduce a cell’s ability to acquire information about its environment. Such information is essential whenever cells need to respond to changing environments (8,9). It is acquired by signaling pathways that modulate the activity or concentration of transcription factors (TFs), which up-regulate or down-regulate effector genes. Thus, reducing gene expression noise can increase the ability of a regulated gene to capture information about a TF’s changing concentration or activity, which is fundamental to produce an optimal cellular response to environmental change (10).

Gene expression regulation is being studied by many researchers whose insights improve our capacity to control and modify living systems (11–14). However, we still do not fully understand how some elementary properties of the interaction of a TF with its binding site on DNA affect the stochastic dynamics of gene expression and the acquisition of information (15). One of these properties is a TF’s residence time on its DNA binding site – the mean time that a TF remains bound to DNA. The residence time is equal to the inverse of the dissociation rate *k*_*d*_ between a TF and DNA. Theoretical and experimental work has shown that the dissociation rate can affect gene expression, affect the size of gene expression bursts (16–19), and modulate gene expression noise (20–23). However, it is difficult to discern whether the dissociation rate affects gene expression by altering the residence time or the affinity between a TF and DNA, because both depend on the dissociation constant *k*_*d*_ (affinity is given by the ratio *K*_*eq*_=*k*_*d*_/*k*_*a*_ [M], where *k*_*a*_ is association rate between a TF and DNA).

Many TFs bind DNA transiently, with residence times ranging between seconds and minutes (24–35). Such TFs include MYC, p53, and glucocorticoid receptors, which are involved in fundamental processes such as apoptosis, DNA repair, DNA maintenance, and stress responses. They also include pioneer TFs that directly interact with chromatin and open it (36–38). The duration of a TF’s residence time on a specific binding site can vary, even for different tissues within the same organisms, by chromatin modifications and the interaction of the TF with other molecular components (24,39,40). Such variation implies that residence time may play a role in regulating gene expression. Nevertheless, we do not know how residence time affects gene expression noise, because of limitations in experimental technology. For example, it is difficult to measure residence time and simultaneously quantify the rate of gene expression. It is also hard to quantify the number of TF binding events at a specific binding site in a given time (16,24,31). Moreover, it is challenging to experimentally modify only residence time without also affecting affinity.

Here we circumvent these limitations through a stochastic model of gene expression regulation. With this model, we study how residence time affects gene expression noise and the amount of information acquired by a gene expression system. Our analyses show that the effect of residence time increases as the level of induction of a gene decreases. At high induction levels, residence time has no effect on gene expression. However, as induction levels decrease, shorter residence times reduce the amount of gene expression noise and produce more regular gene expression dynamics. Shorter residence times also increase a gene regulation system’s capacity to acquire information about the concentration of a TF. In sum, shorter residence times improve a gene’s response to changes in its cellular environment.

## Results

### Model and main concepts

We use a two-state model of gene expression that represents the transcriptional activation and inactivation of a gene. In this model, TF molecules associate and dissociate from the gene’s transcription factor binding site (*DNA*_*bs*_) at rates *k*_*a*_(M^−1^s^−1^) and *k*_*d*_(s^−1^), respectively. The regulated gene is expressed only when its binding site is bound by a TF, in which case the gene is transcribed into mRNA at rate *k*_1_. The resulting mRNA is then translated into protein molecules at rate *k*_2_. Finally, mRNA and protein molecules degrade at rates *d*_1_ and *d*_2_, respectively (Fig. 1a).

**Fig 1.**
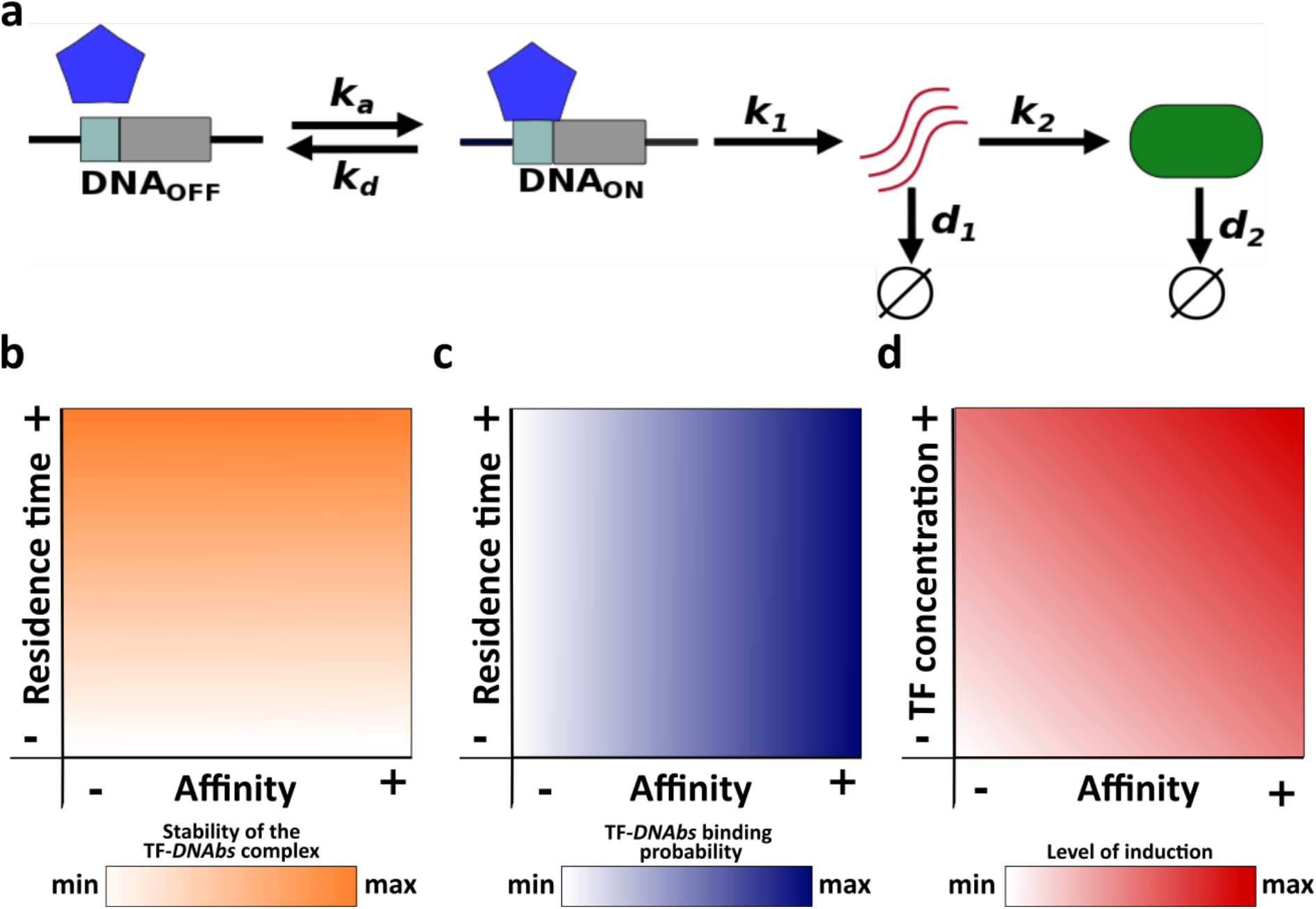
Schematic description of the model and main concepts. (a) *k*_*a*_ and *k*_*d*_ correspond to the association and dissociation rate, respectively; *k*_1_ and *k*_2_ correspond to the mRNA and protein synthesis rate, respectively; *d*_1_ and *d*_2_ correspond to the mRNA and protein degradation rate, respectively. Relationships of both residence time and affinity with (b) the stability of the TF-*DNA*_*bs*_ complex, and (c) TF-*DNA*_*bs*_ binding probability. (d) Relationship of affinity and TF concentration with the level of induction.

Residence time is the average life span or half-life (*t*_1/2_) of the TF-*DNA*_*bs*_ complex. In other words, residence time quantifies the stability of this complex (Fig. 1b). Affinity is quantified with the equilibrium constant *K*_*eq*_. The equilibrium constant is equal to the concentration of free TF at which half of all binding sites are occupied. A high equilibrium constant is equal to a low affinity, because it means that a large concentration of TF is required to occupy 50% of binding sites. For a given TF concentration, the probability that a binding site is occupied increases with increasing affinity (Fig. 1c).

Although both affinity and residence time depend on the dissociation rate *k*_*d*_, they can be modified independently from each other. Changing the dissociation rate will modify the residence time, but it can leave the affinity unchanged if the association rate changes appropriately to keep the ratio *k*_*d*_/*k*_*a*_ constant. Conversely, by changing only the association rate, affinity can be modified without altering residence time.

Notice that affinity and TF concentration jointly determine the level of induction of gene expression, because a gene is more likely to be active when the concentration of free TF is higher than its affinity to DNA. Conversely, when this concentration is lower than the affinity, the gene will tend to be inactive. One can increase the level of a gene’s induction by increasing either TF concentration or affinity (Fig. 1d). Below we change the level of induction by modifying TF concentration, but changing affinity itself yields the same observations (see Sup Inf 1).

We simulate gene regulation dynamics using Gillespie’s algorithm (41), which reproduces the stochastic dynamics of many chemical systems, using biologically meaningful values of all biochemical parameters (Sup Table 1 and Methods). Because both the mRNA and protein output of our modeled gene regulation system behave qualitatively identically, we focus on the protein output below (see Sup Figs for mRNA).

### Short residence times reduce gene expression noise and modulate gene expression dynamic

We first study the effect of residence time and affinity on gene expression noise. To quantify noise, we quantified the size of the temporal fluctuations in the number of proteins, as the difference between the maximal and the minimal number of expressed protein molecules 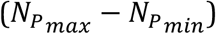, and averaged this difference over 1000 simulations. Two alternative noise measures, the coefficient of variation and the Fano factor yield identical observations (see Sup Inf 2 and Sup Fig 1).

In these simulations, we varied residence time within the interval [1s,1h], TF concentration within the interval [10^−11^M,10^−*7*^M], and set the affinity to 10^−9^M. Notice that the TF concentration interval ranges two orders of magnitude below and above the affinity, which implies that the level of gene induction ranges from almost always inactive to almost always active. Hence, high and low TF concentration values correspond to high and low induction levels, respectively.

At the highest TF concentration, residence time does not affect noise (Fig. 2a; Sup Fig. 2a). However, as the TF concentration decreases, a longer residence time increases noise (Fig. 2a; Sup Fig. 2a). For example, individual protein expression trajectories at extremely short (*t*_1/2_=1s) and long (*t*_1/2_=1h) residence times are very similar at the highest TF concentration (Fig. 2b; Sup Fig. 2b). However, as TF concentration decreases, long residence times cause sporadic, but large fluctuations in protein expression (Fig. 2c and d; Sup Fig. 2c and d).

**Fig 2.**
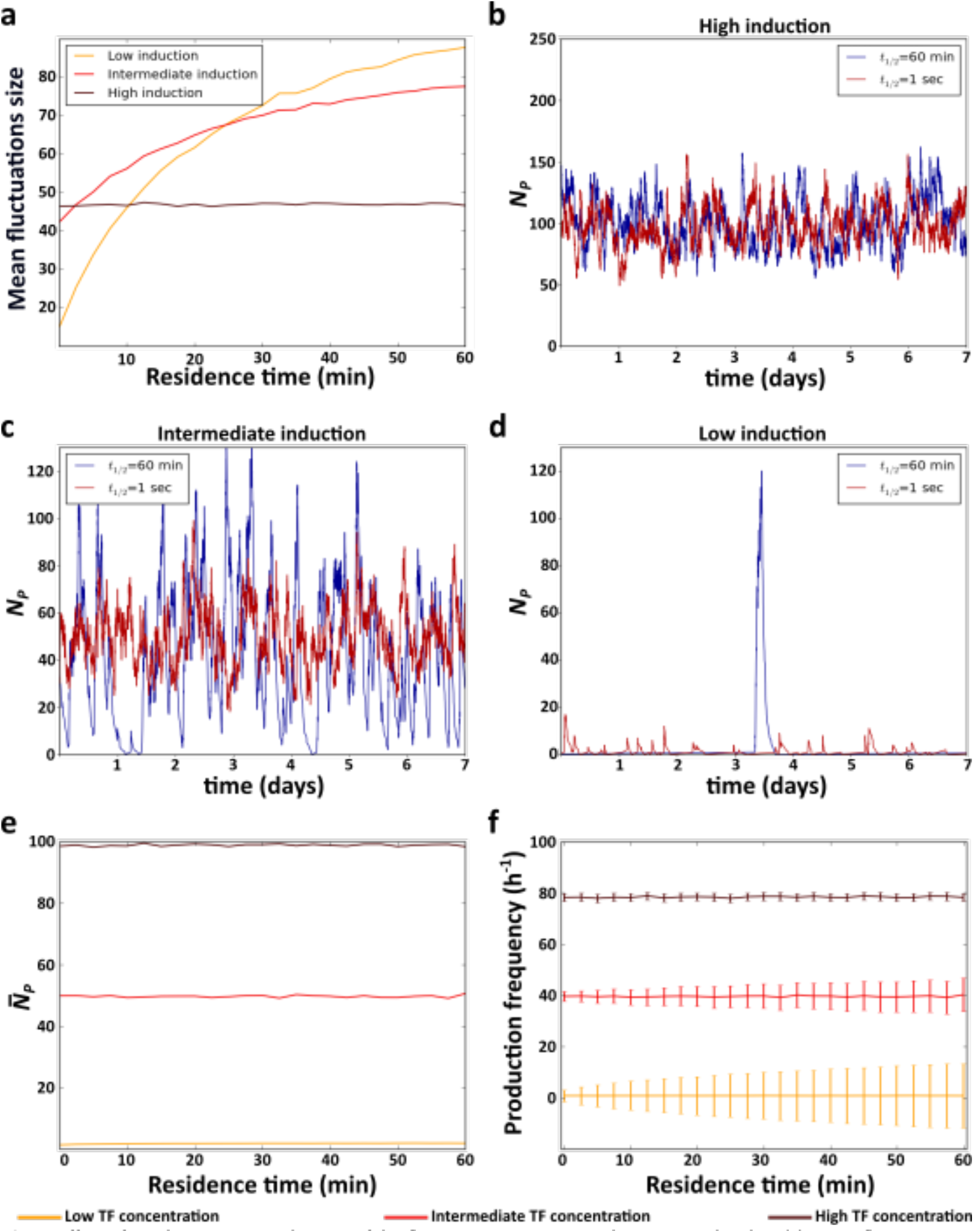
Effect of residence time on the size of the fluctuations in expressed protein molecules. (a) Mean fluctuation size in the number of protein molecules (*y* axis) at different levels of induction as a function of residence time (*x* axis). (b-d) Example time trajectories of the number of protein molecules *N*_*P*_ at three different levels of induction. Analyses of (e) mean number of protein 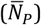 (f) mean and coefficient of variation of the frequency of protein production events. (b-d) Red and blue lines show data for short (1s) and a long (1h) residence times, respectively. (a, e and f) High (TF=10^−7^M), intermediate (TF=10^−9^M) and low (TF=10^−11^M) levels of induction are indicated in the color legend below the figure.

To understand these observations, notice that when the TF concentration is very high, the level of induction is high because a TF molecule is bound to DNA most of the time. In this case, gene expression resembles that of a constitutive gene, regardless of the TF’s residence time. In consequence, gene expression noise is only determined by the degradation and synthesis rates of mRNA and protein (1,2). In other words, it is independent of residence time (Fig. 2a and b; Sup Fig. 2a and d).

This is no longer true as the TF concentration decreases. In this case, the probability that a TF is bound to DNA at any one time decreases, and longer residence times increase the average amount of time that a TF is either bound or unbound. In other words, longer residence times produce longer periods of active and inactive gene expression. During active periods, proteins are produced, whereas during inactive periods, previously expressed proteins decay. Thus, longer residence times lengthen both active and inactive periods, which results in large fluctuations in the number of proteins (Fig. 2c and d; Sup Fig. 2c and d). Reducing the residence time (at constant induction) decreases the duration of both active and inactive periods by the same amounts. As a result, expressed molecules accumulate and decay for shorter time periods, and fluctuations in these molecules become smaller (Fig. 2a–d; Sup Fig. 2a−d).

In contrast to its effects on noise, residence time does not affect the mean level of protein expression, which only depends on the level of induction (Fig. 2e; Sup Fig. 2e). The reason is that the frequency of both protein production and degradation events (i.e., the mean number of proteins produced and degraded in a given period of time) is not affected by residence time, regardless of the level of induction (Fig. 2f; Sup Fig. 2f and 3a and b). However, as the level of induction decreases, shorter residence times decrease variation in the frequency of production and degradation events (Fig. 2f; Sup Fig. 2f and 3a and b). As a consequence, protein expression is more homogeneous at shorter residence times, because protein production and degradation events alternate more regularly, except at the very lowest affinities (Sup Inf 3 and Sup Fig. 3c-f).

**Fig 3.**
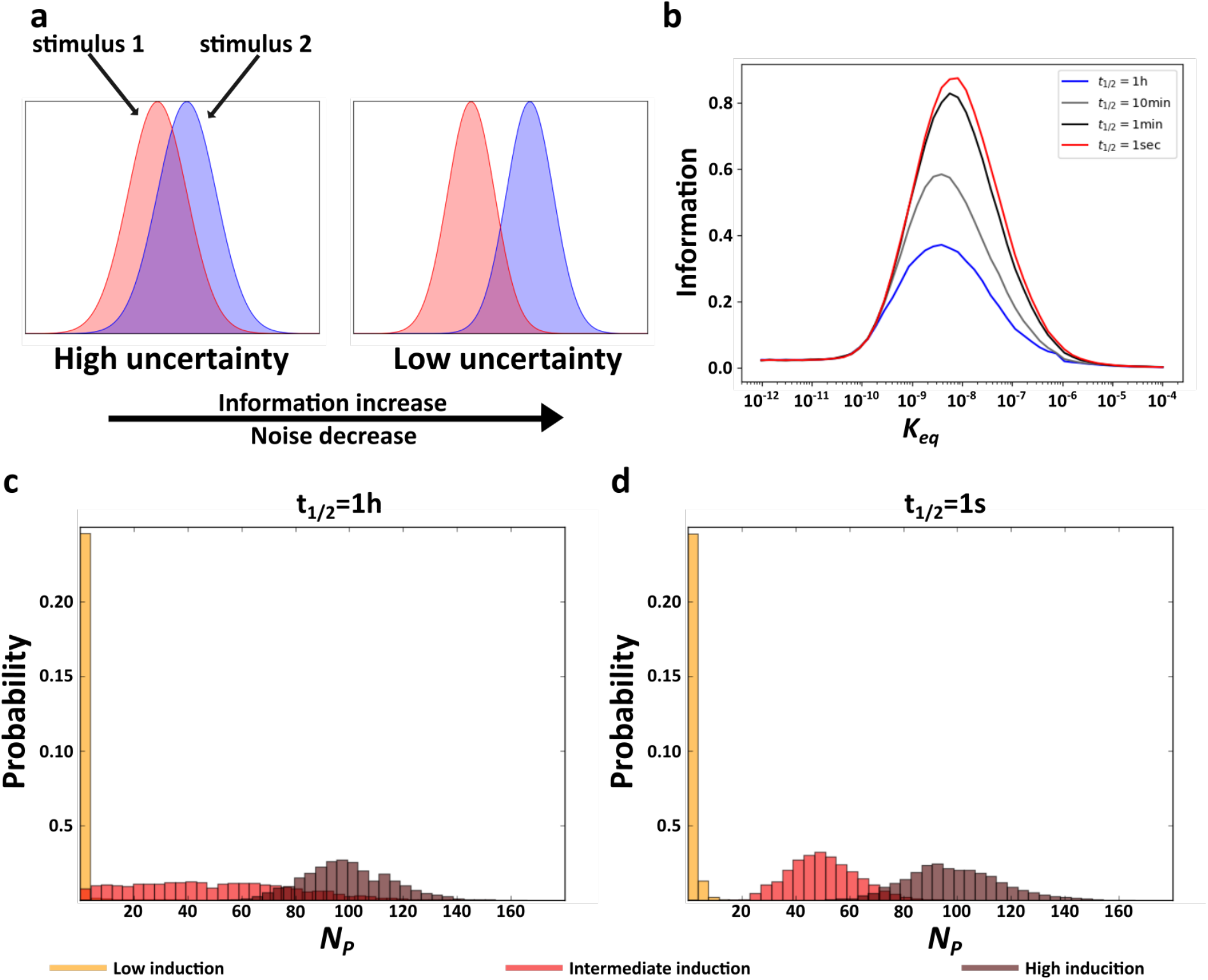
Residence time and information. (a) Schematic explanation of the relationship between noise and acquired information. The panel shows hypothetical response protein distributions produced by two different stimuli at a high (right) and a low (left) level of gene expression noise. (b) Acquired information at different residence times as a function of the affinity (*K*_*eq*_) between TF molecules and *DNA*. (c and d) Distributions of the number of expressed proteins *N*_*P*_ at three different TF concentrations (see color legend) with a long (c), and a short (d) residence time. In c and d, *K*_*eq*_ =10^−9^M.

### Residence time and information

Due to noise, the regulation of gene expression transforms a concentration of a TF into a distribution of expressed mRNA and protein molecules. Different TF concentrations may produce overlapping distributions of expressed molecules, in which case information about TF concentration gets lost. Reducing gene expression noise can reduce this overlap and thus also the amount of lost information. (Fig. 3a) (10). Because shorter residence times reduce gene expression noise (Fig. 2), we hypothesized that they also increase the amount of information protein which expression levels contain about the concentration of the regulating TF. To find out, we quantified the mutual information between protein expression and TF concentration. Mutual information is an information theoretical quantity that encapsulates the reduction in uncertainty about one random variable provided by knowledge about another random variable (42) (see Methods). To quantify mutual information, we performed 2500 stochastic simulations of gene expression dynamics for each of *n* evenly distributed TF concentrations within the interval [10^−7^M/*n*,10^−7^M], exploring affinity values within the interval [10^−12^M,10^−4^M]. This range includes affinity values below the minimal TF concentration, where induction is low regardless of TF concentration, and above the maximal concentration, where induction is high regardless of TF concentration.

In earlier work, we have shown that the amount of information that gene expression levels contain about the concentration of the regulating TF depends on the affinity between a TF and its binding site. At very low affinities, gene regulation is insensitive to TF concentration, such that gene expression conveys little information. At very high affinities, the level of gene induction is high for all TF concentrations, such that gene expression also conveys little information – it is similar to that of a constitutively expressed gene for all TF concentrations (43). Our current work shows that this pattern holds regardless of residence time (Fig. 3b; Sup. Fig. 4a).

In contrast, residence time does affect acquired information at intermediate affinities. Specifically, even though the mean number of expressed molecules does not depend on a TF’s residence time on DNA (Fig. 2e; Sup. Fig. 2e), their variability decreases with shorter residence times (Fig. 2a). As a result, as residence time decreases, the overlap between protein distributions decrease (Fig. 3c and d; Sup Fig. 4b and c), which increases the amount of acquired information (Fig. 3b; Sup Fig. 4a). In sum, under conditions where a gene regulation system can acquire information, shortening residence time increases the amount of acquired information.

## Discussion

Previous theoretical and experimental work showed that gene expression noise can be modulated by the dissociation rate *k*_*d*_ of a DNA-bound TF (16–23), but this work did not distinguish between the effects of residence time and affinity. This distinction, however, is important because both properties depend on the dissociation rate but have different effects on gene expression dynamics. Here, we study these properties separately, and show that short residence time can reduce expression noise, while in general noise increases as affinity decreases (Fig 2).

Our results also show that the effect of residence time on gene expression is not independent from affinity. When a gene is highly induced, residence time does not affect gene expression. However, as the level of induction decreases, short residence times can help produce less noisy expression. Short residence times effectively fragment gene activity into short periods of active and inactive expression, which prevents the excessive accumulation and depletion of proteins, and thus also excessive stochastic variation in gene expression. Similarly, the effect of affinity on noise depends on residence time. In particular, the effect of affinity is reduced when residence time decreases. However, noise can even decrease with decreasing affinity when residence times are very short and in the range of seconds (Fig. 2a).

In previous work, we showed that expressed proteins harbor information about the concentration of a TF regulating their expression, if the TF’s affinity to regulatory DNA is of the same order of magnitude as its concentration (43). Here, we show that this behavior is independent of residence time. However, because shorter residence times render gene expression less noisy, the overlap between protein expression distributions resulting from different TF concentrations decreases. Consequently, shorter residence times increase the ability of a gene regulation system to distinguish between different TF concentrations.

Recent technological advances will permit experimental verification of our observations. First, in the last decade methods to quantify the ability of a regulated gene to acquire information about the concentration of a TF have become standardized (10,44). Also, the effect of residence time on noise can be quantified with techniques that simultaneously quantify TF-DNA binding events and the production of mRNA molecules, which are being developed (16). It is especially difficult to obtain multiple alleles of a TF with different residence times without also altering the TF’s affinity to DNA. However, because our model shows that affinity affects gene expression by changing the induction level at a specific TF concentration, our observations hold regardless of whether one varies TF concentration or affinity (Sup Inf 1). Hence, to test the effect of residence time on noise and information acquisition, one can compensate for any change in affinity by adjusting a gene’s induction level with TF concentration.

Our two state model of gene regulation is simple and does not represent processes such as the binding of RNA polymerases to DNA explicitly (45). However, previous work has shown that the two state model produces similar gene expression dynamics as more complex models (19), suggesting that our main results may hold for such models. Nevertheless, each additional regulatory requires time, which may constrain the amount of time that a TF must be minimally bound to DNA before it can affect gene expression (46). Future work also needs to consider other kinetic parameters regulating gene expression, such as the mRNA synthesis rate, because noise and information can be affected by these parameters (1,2,20,22,47–49).

Our results are in agreement with previous work and complement this work. For example, experimental evidence suggests that the affinity of essential regulators of gene expression, such as NF-κB and TBP, modulates gene expression noise (50,51). Moreover, a model based on the binding dynamics of Sox2 and Oct4, two important regulators of the pluripotency of stems cells, showed that long residence times reduce the sensitivity of gene expression to TF concentration, because TFs with long residence times are bound to DNA most of the time regardless of their concentration (26,31). Another study showed that negative regulatory feedback loops in general suppress noise more effectively when residence times are short (23).

Our work also helps solve an apparent contradiction between experimental and theoretical work about the importance of residence time. In particular, it has been predicted that longer residence times facilitate gene expression, because they increase the probability of a successful activation of gene expression by TFs, by providing longer time for other components, such as polymerases, to successfully bind DNA (24,25,52–54). Experimentally measured residence times, which are generally short and lie within seconds to minutes (24–35), are inconsistent with this prediction. Our work shows that residence time does not affect average gene expression levels. Instead, residence time reduces expression noise and can help signaling systems acquire information without modifying the probability of a successful activation of gene expression, which can help explain why short residence times may be prevalent in nature.

## Methods

### Two-state model of gene expression

To study how a transcription factor’s (TF’s) residence time on DNA affects gene expression, we built a gene expression model in which a *TF* binds to a DNA binding site (*DNA*_*bs*_) to regulate the expression of a nearby gene. *TF* molecules associate with the DNA binding site at a rate *k*_*a*_ (M^−1^s^−1^), and dissociate from it at a rate *k*_*d*_ (s^−1^). Only when the TF is bound to DNA does transcription occur (at a rate *k*_1_ [s^−1^]). Transcribed mRNA molecules are degraded at a rate *d*_1_ (s^−1^). Proteins are translated from mRNA molecules at a rate *k*_2_ (s^−1^), and degraded at a rate *d*_2_ (s^−1^).

### Stochastic simulations

To simulate the behavior of our gene expression model, we use Gillespie’s discrete stochastic simulation algorithm (41), using the numpy python package for scientific computing (http://www.numpy.org/). Gillespie’s algorithm captures the stochastic nature of chemical systems. It assumes a well-stirred and thermally equilibrated system with constant volume and temperature. The algorithm requires the probability *p*_*j*_ that a chemical reaction *R*_*j*_ occurs in a given time interval [t,t+τ). This probability *p*_*j*_ is proportional to both the reaction rate and the number of reacting molecules. For the reversible bindings of TF molecules to DNA, the association probability *p*_*a*_ and the dissociation probability *p*_*d*_ are given by

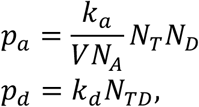

where *V* is the reaction volume, *N*_*A*_ is Avogadro’s number, and *N*_*T*_, *N*_*D*_, and *N*_*TD*_ are the numbers of TF molecules, DNA binding sites, and TF-*DNA*_*bs*_ complexes. Notice that the dissociation of TF-*DNA*_*bs*_ complexes is a first-order reaction, which is independent of the volume in which the reaction takes place. In contrast, the association of TF molecules with DNA binding sites is a second-order reaction, which is inversely proportional to the volume.

The probabilities *p*_*mRs*_, *p*_*mRd*_, *p*_*Ps*_ and *p*_*Pd*_ of mRNA transcription, mRNA degradation, protein synthesis, and protein degradation are given by

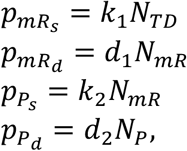

respectively. In these expressions, the quantities *N*_*TD*_, *N*_*mR*_ and *N*_*P*_ are the numbers of TF-DNA complexes, mRNA molecules and of protein molecules, respectively. Because we model a haploid organism with only a single non-leaky DNA binding site, the probability of mRNA synthesis can be reduced to

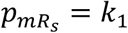

when the DNA is bound by the transcription factor (*N*_*TD*_=1), and to

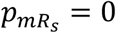

when it is unbound (*N*_*TD*_=0).

### Initial conditions for simulations

We assume that the TF a concentration of 10^−7^M corresponds to a few thousand TF molecules per cell, a realistic number in animal and yeast cells (55,56). Because we model only one binding site, the concentration of free TF is not substantially affected by the binding of a single TF molecule to *DNA*. We therefore do not distinguish between the free and the total TF concentration. After this simplification, to determine the initial conditions of our model, we calculate the probability 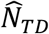 that the binding site is bound by a TF molecule as

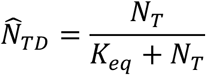

where *N*_*T*_ is the total number of TF molecules. We selected the initial state of the DNA 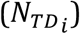 at random with binomial probability 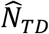 (i.e., 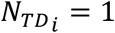 if the DNA is bound by a TF molecule, and zero otherwise. It follows that

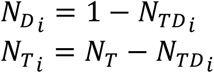

where 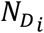 is the initial state of the number of non-bound DNA binding site and 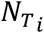 is the number of free TF molecules. As the initial state of the number mRNA and protein molecules we used

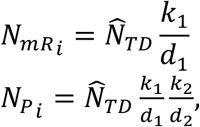

which are the expected average number of mRNA and protein molecules for a constitutively expressed gene (2), multiplied by the probability that the DNA is bound by a TF molecule.

### Information quantification

The number of molecules of any chemical species in a cell or in a unit volume fluctuates, because molecules are produced and decay stochastically. We use Shannon’s entropy to quantify the unpredictability caused by such stochastic fluctuations in the number of transcription factor molecules as

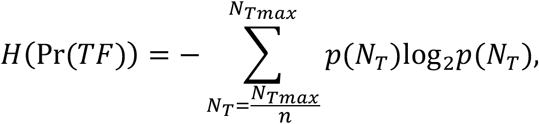

where Pr(*TF*) is the distribution of the number of transcription factor molecules (*N*_*T*_), and *p(N*_*T*_) is the probability that the system contains *N* molecules of the transcription factor.

To estimate information we performed simulations from *n* different numbers of transcription factor molecules that were evenly distributed within the interval [*N*_*Tmax*_/*n*,*N*_*Tmax*_] (*n*<*N*_*Tmax*_). For this reason

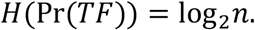

TF-DNA binding triggers the transcription of mRNA molecules that are then translated into protein molecules. We use the number of mRNA *N*_*mRNA*_ and protein molecules *N*_*P*_ as the system’s response or output, which we denote as *O*.

A gene expression system acquires information when the number of expressed proteins or mRNA reflects the number of transcription factors. This information can be quantified via the mutual information

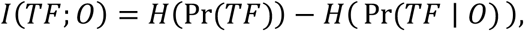

a widely used quantity in information theory (42). It is equal to the difference between the entropy *H*(Pr(*TF*)) and the conditional entropy *H*(Pr(*TF*|*O*)), which represents the entropy of the number of transcription factor molecules for a given number of mRNA or protein molecules. In other words, the mutual information *I* quantifies the amount of information that the number of expressed mRNA or protein molecules harbors about the number of transcription factor molecules.

### Noise quantification

The model produces a probability distribution of the number of mRNA and protein molecules for any given number of transcription factor molecules *N*_*T*_. This response is thus a conditional probability distribution, which we write as

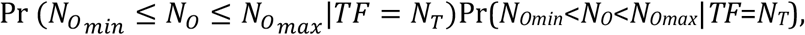

where *N*_*Omin*_ and *N*_*Omax*_ are the minimal and maximal number of mRNA or protein molecules, respectively. We performed 1000 simulations to estimate noise using three different measures. The size of the fluctuation, Fano factor and the coefficient of variation. The size of the fluctuations was quantified as the average difference 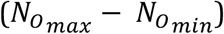. Fano factor as the variance of the response distribution divided by its mean 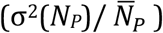. Coefficient of variation as the standard deviation of the response distribution divided by its mean 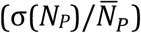.

### Parameter values

Our simulations considered biologically sensible parameter ranges. Specifically, for TF-*DNA*_*bs*_ binding, empirical data suggests that usually *K*_*eq*_<10^−8^ and can reach picomolar (10^−12^M) or even smaller values (46,55,57–60). Because experimental research has shown that residence time varies from seconds to tens of minutes (24,27,29), we used dissociation rates producing residence time within this interval [1s,1h].

For mRNA, experimentally measured half-lives usually lie in the range of seconds to hours (61–64). Protein half-lives are usually longer than mRNA half-lives (65) and lie between hours and days (63,66). Taking all this information into consideration we chose mRNA half-lives of ~3.3min, and protein half-lives of ~1.5h. We assumed that the ratio *k*_2_/*k*_1_ of the protein synthesis rate to the mRNA synthesis rate exceeds 1.0 (67).

---

**Sup Fig 1.**
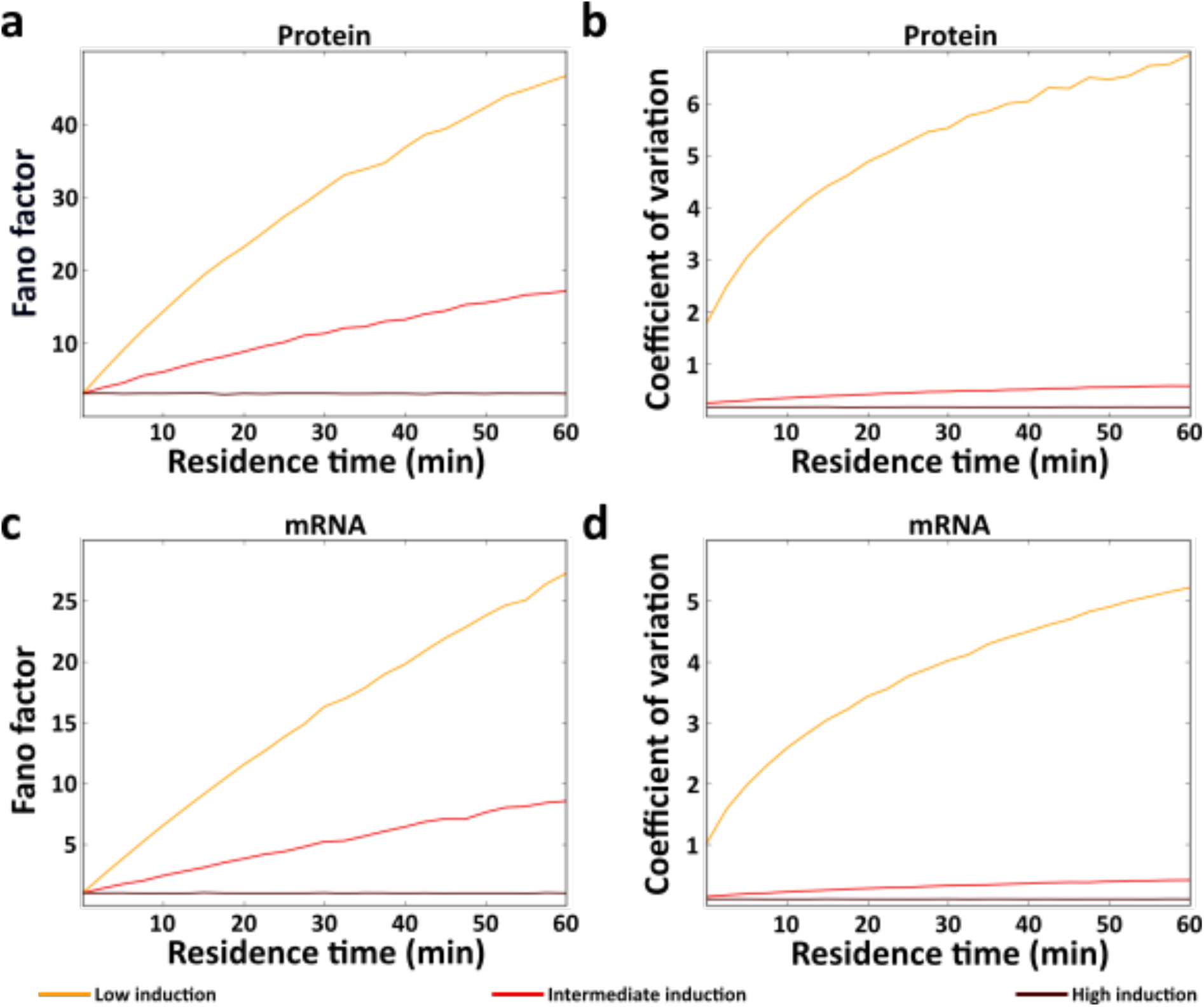
Effect of residence time on Fano factor and coefficient of variation. (a and c) Fano factor and (b and d) coefficient of variation in the number of expressed protein (a and b) and mRNA (c and d) molecules as a function of residence time (*x* axis) at a high (TF=10^−7^M), intermediate (TF=10^−9^M) and low (TF=10^−11^M) levels of induction, as indicated by the color legend below the figure.

**Sup Fig 2.**
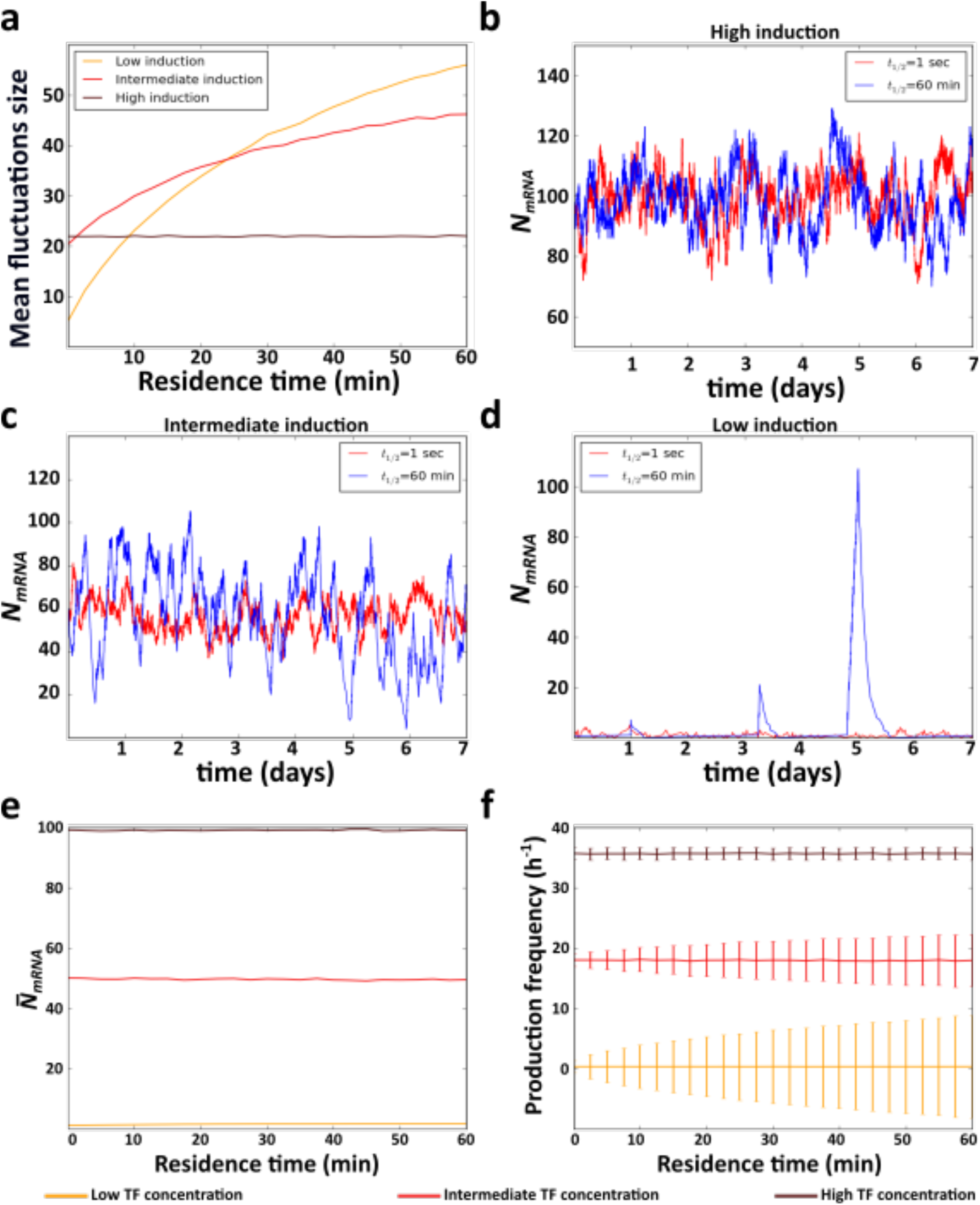
Effect of residence time on the size of the mRNA fluctuations. (a) Mean fluctuation size in the number of mRNA molecules (*y* axis) at a high (TF=10^−7^M), intermediate (TF=10^−9^M) and low (TF=10^−11^M) level of induction, as a function of residence time (*x* axis). (b-d) Example time trajectories of the number of expressed mRNA molecules *N*_*mRNA*_ obtained from the simulation of the model at three different levels of induction. Red and blue lines show data for short (1s) and long (1h) residence times, respectively. Analyses of (e) mean number of mRNA 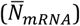 (f) mean and coefficient of variation of the frequency of mRNA production events at high (TF=10^−7^M), intermediate (TF=10^−9^M), and low (TF=10^−11^M) TF concentrations, as indicated in the color legend below the figure.

**Sup Fig 3.**
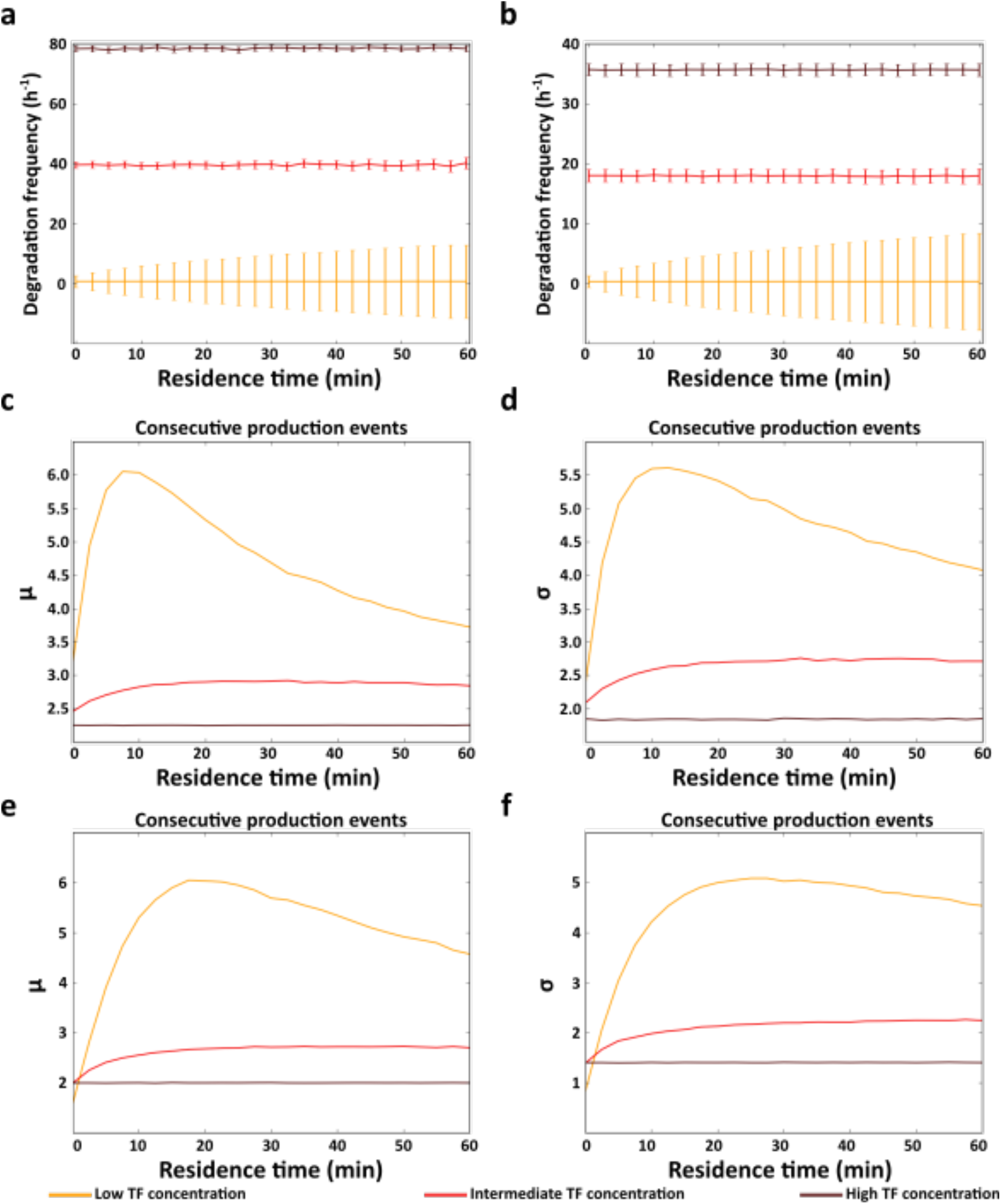
Residence time and both production and degradation events. Mean and standard deviation of (a) protein and (b) mRNA degradation events. (c and e) Mean and (d and f) standard deviation of the number of consecutive (c and d) protein and (e and f) mRNA production events as a function of residence time (*x* axes). All analyses were performed at high (TF=10^−7^M), intermediate (TF=10^−9^M) and low (TF=10^−11^M) affinity values (see color legends)

**Sup Fig 4.**
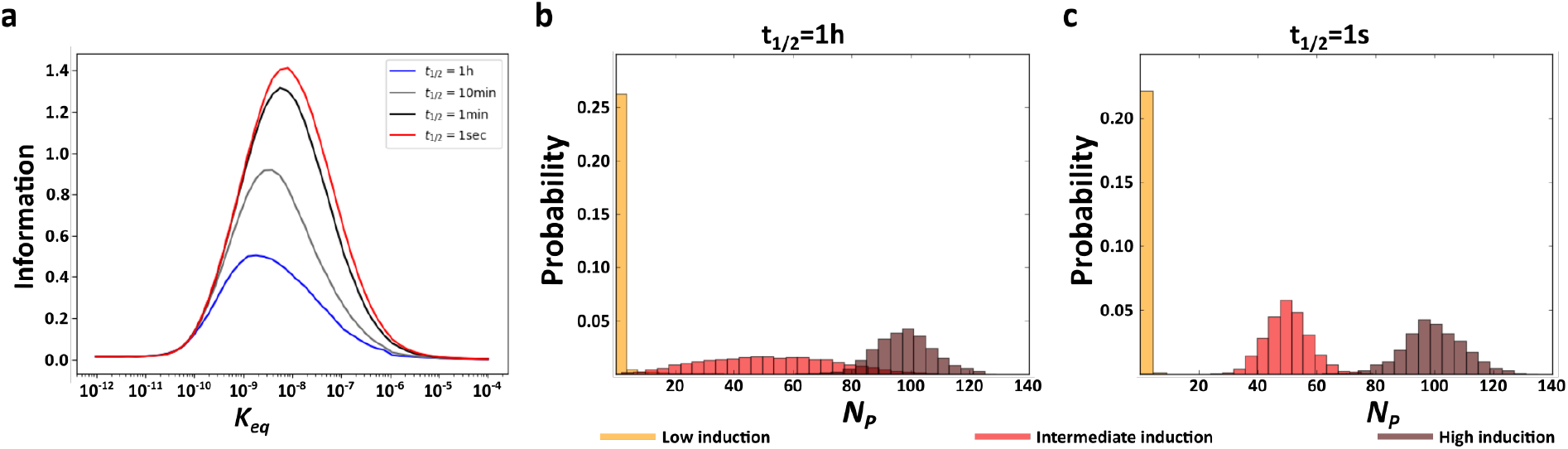
Residence time and acquired information quantified through mRNA expression. (a) Information acquired at different residence times as a function of the affinity (*K*_*eq*_) between TF molecules and *DNA*. (b and c) Distributions of the number of proteins *N*_*P*_ produced at three different TF concentrations with a long (b), and a short (c) residence time. In c and d, *K*_*eq*_ =10^−9^M.

**Sup Fig 5.**
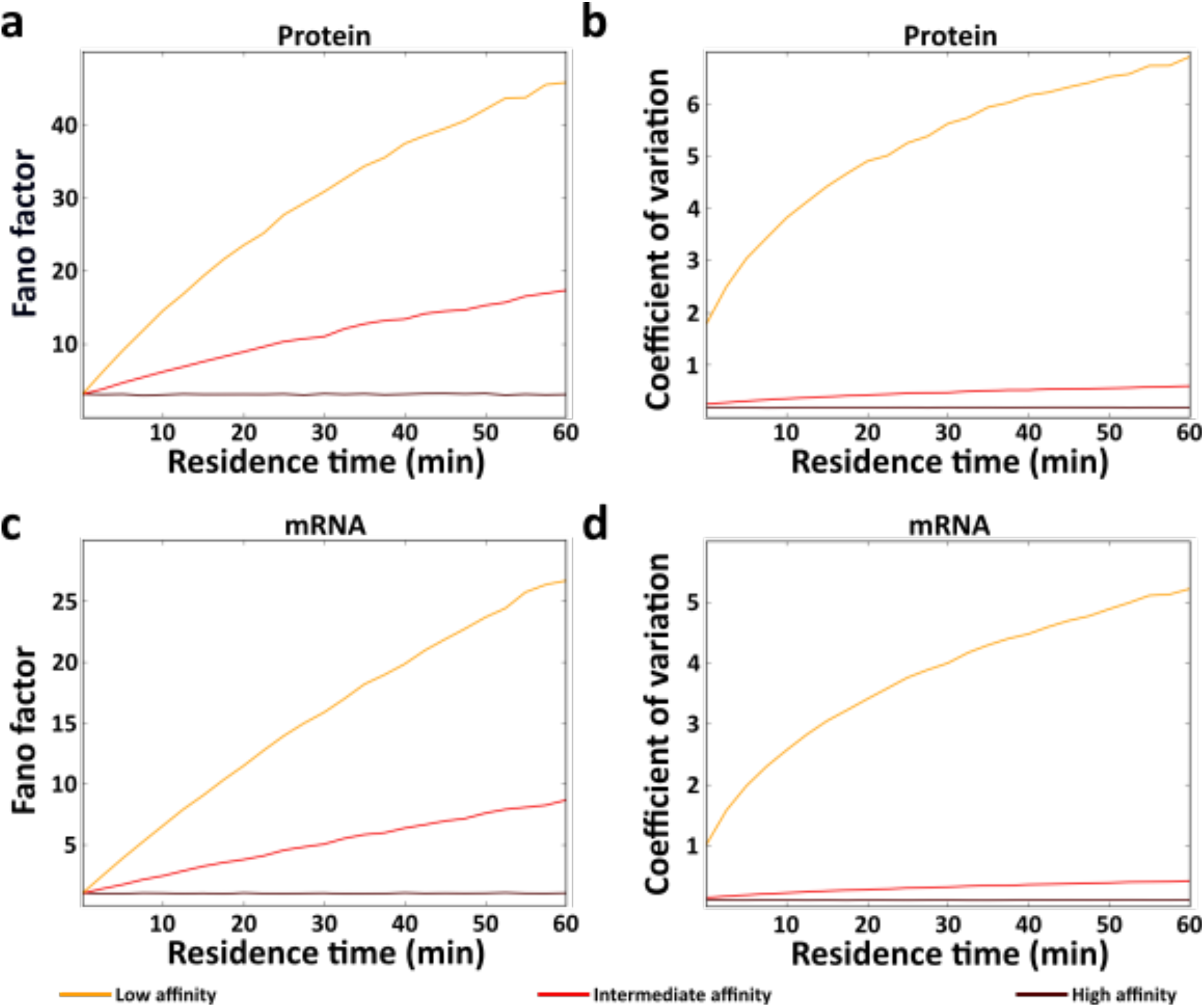
Effect of residence time on Fano factor and coefficient of variation. (a and c) Fano factor and (b and d) coefficient of variation in the number of protein (a and b) and mRNA (c and d) molecules as a function of residence time (*x* axes) at high (*K*_*eq*_=10^−11^M), intermediate (*K*_*eq*_=10^−9^M) and low (*K*_*eq*_=10^−7^M) affinity values, as indicated by the color legend below the figure.

**Sup Fig 6.**
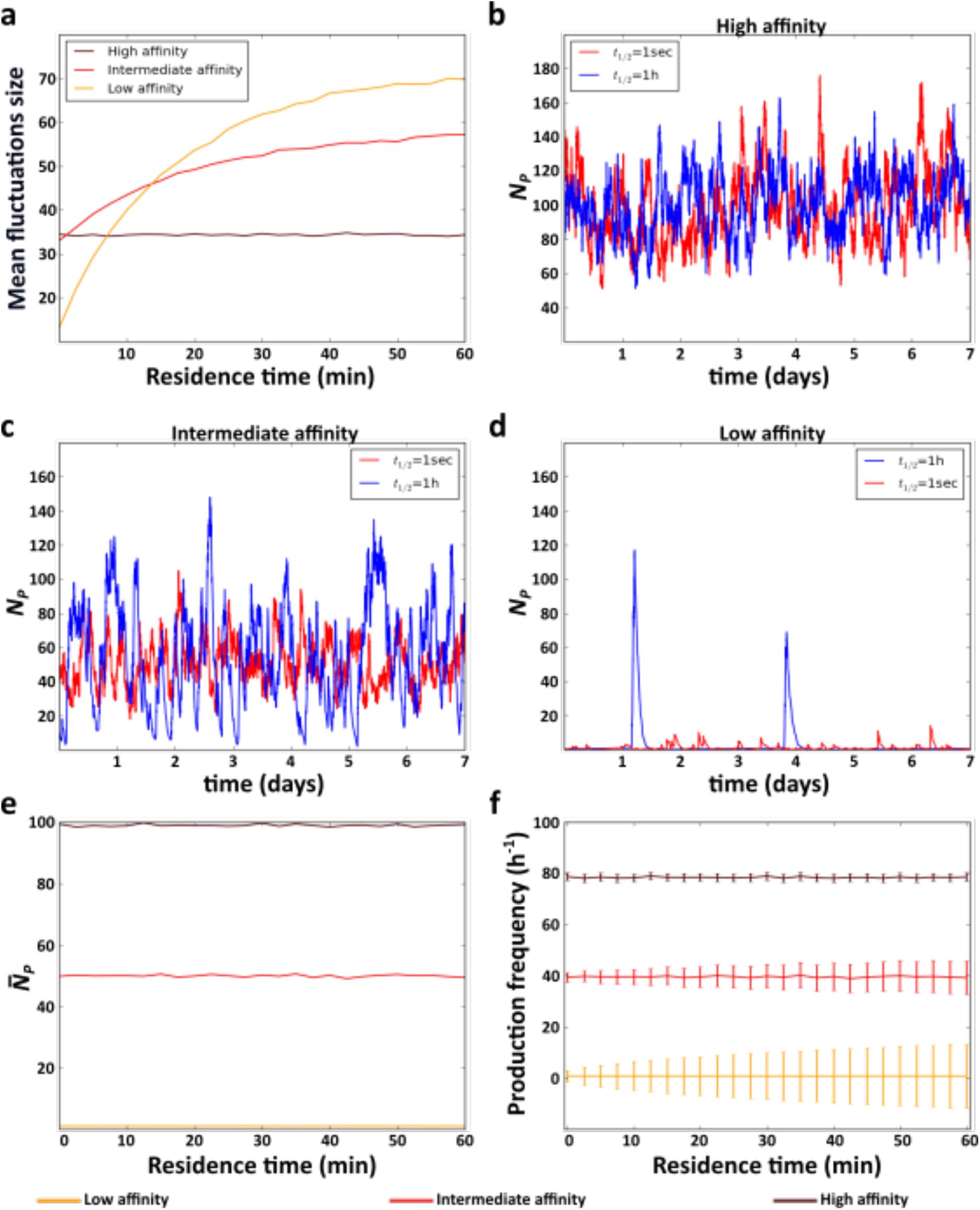
Effect of residence time on the expression dynamic of protein molecules. (a) Mean fluctuation size in the number of protein molecules (*y* axis) at high (*K*_*eq*_=10^−11^M), intermediate (*K*_*eq*_=10^−9^M) and low (*K*_*eq*_=10^−7^M) affinities, as a function of residence time (*x* axis). (b-d) Example time trajectories of the number of protein molecules *N*_*P*_ obtained from the simulation of the model at the three different affinities. Red and blue lines show data for short (1s) and long (1h) residence times, respectively. Analyses of (e) mean number of proteins 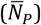, and (f) mean and coefficient of variation of the frequency of protein production events.

**Sup Fig 7.**
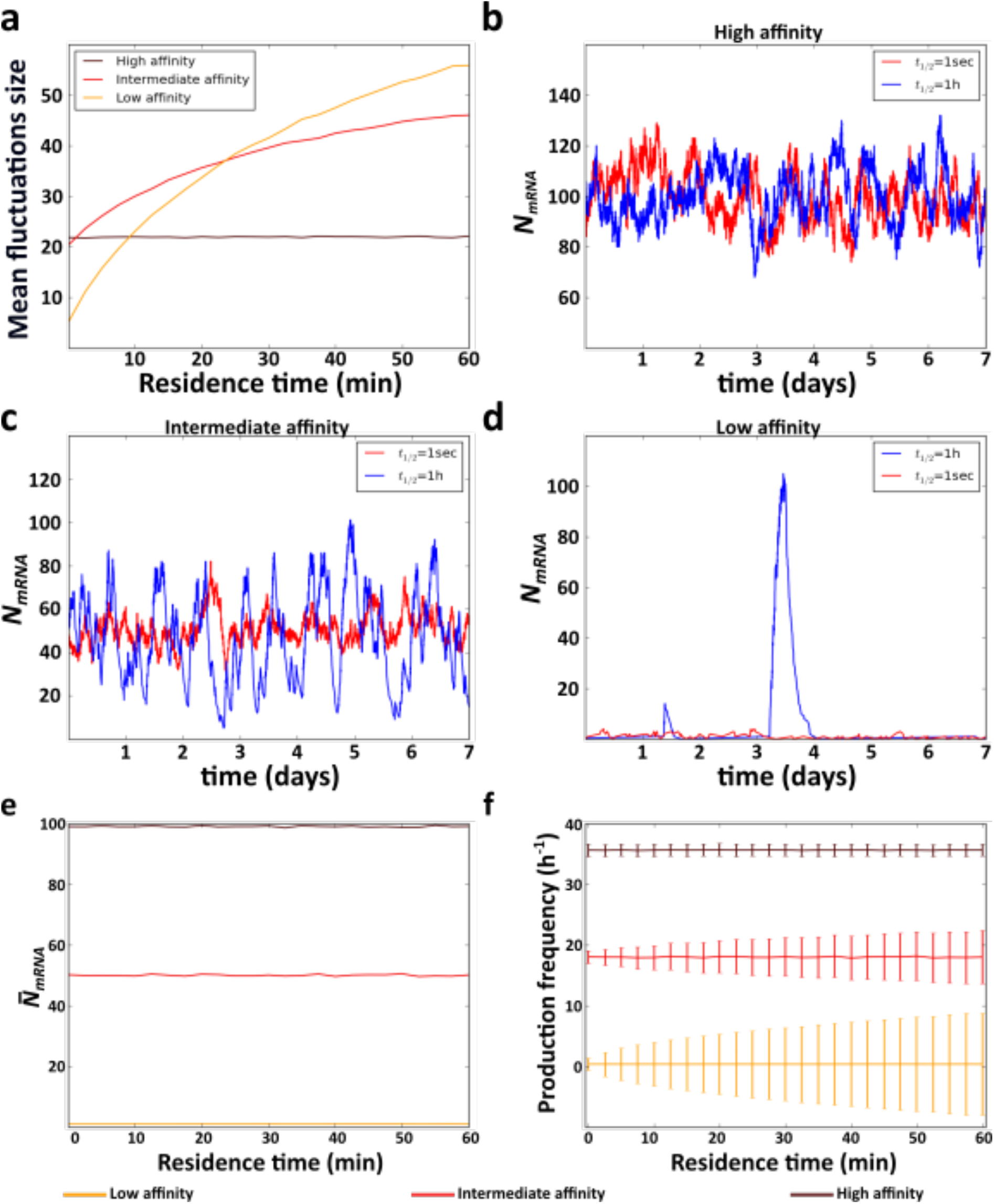
Effect of residence time on time on the expression dynamic of mRNA molecules. (a) Mean fluctuation size in the number of mRNA molecules (*y* axis) at high (*K*_*eq*_=10^−11^M), intermediate (*K*_*eq*_=10^−9^M) and low (*K*_*eq*_=10^−7^M) affinity values, as a function of residence time (*x* axis). (b-d) Examples time trajectories of the number of expressed mRNA molecules *N*_*mRNA*_ at three different affinity values. Red and blue lines show data for short (1s) and long (1h) residence times, respectively. Analyses of (e) mean number of expressed mRNA molecules 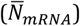, and (f) mean and coefficient of variation of the frequency of mRNA production events

**Sup Fig 8.**
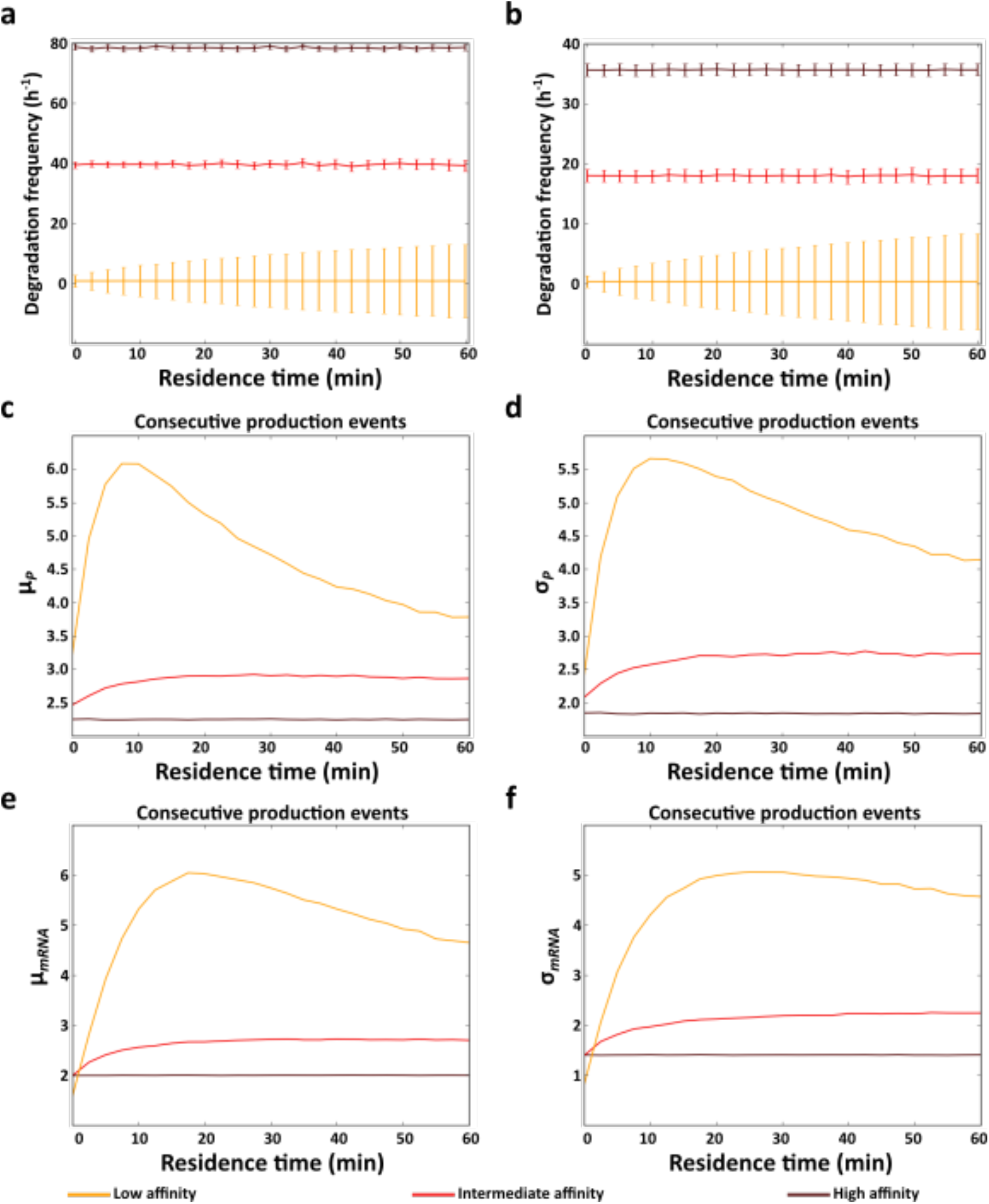
Residence time and both production and degradation events. Mean and standard deviation of (a) protein and (b) mRNA degradation events. (c and e) Mean and (d and f) standard deviation of the number of consecutive (c and d) protein and (e and f) mRNA production events as a function of residence time (*x* axes). All analyses were performed at high (*K*_*eq*_=10^−11^M), intermediate (*K*_*eq*_=10^−9^M) and low (*K*_*eq*_=10^−7^M) affinity values (see color legends)

**Sup Table 1.** Parameter values used for the simulations. *k*_*d*_, *k*_1_, *k*_2_, *d*_1_, *d*_2_ units are s^−1^. *k*_*a*_ units are M^−1^s^−1^. The concentration of the TF is in M.

| | $k_a$ | $k_d$ | $k_1$ | $k_2$ | $d_1$ | $d_2$ | TF conc |
| --- | --- | --- | --- | --- | --- | --- | --- |
| Fig 2<br>Sup Fig 1a | [$2.7 \cdot 10^5$, $10^8$] | [$2.7 \cdot 10^{-4}$, 1] | 0.01 | 0.011 | 0.005 | 0.00022 | [$10^{-11}$, $10^{-7}$] |
| and b<br>Sup Fig 3a,<br>c and d |  |  |  |  |  |  |  |
| Fig 3 | [2.7, 1.0<sup>12</sup>] | [2.7*10<sup>-4</sup>,1] | 0.01 | 0.011 | 0.005 | 0.00022 | [10<sup>-11</sup>,10<sup>-7</sup>] |
| Sup Fig 1c<br>and d<br>Sup Fig 2<br>Sup Fig 3b,<br>e and f | [2.7*10<sup>5</sup>, 10<sup>8</sup>] | [2.7*10<sup>-4</sup>,1] | 0.01 | NA | 0.0001 | NA | [10<sup>-11</sup>,10<sup>-7</sup>] |
| Sup. Fig 4 | [2.7, 1.0<sup>12</sup>] | [2.7*10<sup>-4</sup>,1] | 0.01 | NA | 0.0001 | NA | [10<sup>-11</sup>,10<sup>-7</sup>] |
| Sup Fig 5a<br>and b<br>Sup Fig 6<br>Sup Fig 8a,<br>c and d | [2.7*10<sup>3</sup>, 10<sup>10</sup>] | [2.7*10<sup>-4</sup>,1] | 0.01 | 0.011 | 0.005 | 0.00022 | 10<sup>-9</sup> |
| Sup Fig 5c<br>and d<br>Sup Fig 7<br>Sup Fig 8b,<br>e and f | [2.7*10<sup>3</sup>, 10<sup>10</sup>] | [2.7*10<sup>-4</sup>,1] | 0.01 | NA | 0.0001 | NA | 10<sup>-9</sup> |

## Supplementary information

### 1. Affinity and induction level

As explained in the main text, the affinity of a TF to its DNA binding site and the TF concentration determine the induction level of a gene (Fig 1d). Hence, we hypothesized that the effect of affinity on gene expression could be reproduced by a change in the TF concentration. To find out, we repeated all our analyses, but instead of varying the affinity, we set it to a constant value of 10^−9^M. We then varied the TF concentration within the interval [10^−7^M,10^−11^M]. Analogous to our analysis in the main text, this interval includes TF concentrations two orders of magnitude above and below the affinity. Hence, at the highest TF concentration, the level of induction is high and the regulated gene is almost always active. Conversely, at the lowest TF concentration, the level of induction is low and the regulated gene is almost always inactive. We were able to reproduce all observations we had made by varying affinity through the variation of TF concentrations. Most importantly, as the level of induction decreases, shorter residence times reduce noise (compare Fig 2, Sup Fig. 1 and Sup Fig 2 with Sup Fig. 5-7), producing a more homogenous and regular dynamic of gene expression (compare Sup Fig. 3 with Sup Fig. 8). Hence, our observations show that affinity affects gene expression dynamics mainly through its effect on gene induction.

### 2. Residence time and strength of noise

To study the effect of residence time and affinity on the level of gene expression noise, we also quantified the Fano factor, and the coefficient of variation of gene expression. The Fano factor is equal to the variance in the number of expressed molecules divided by its mean 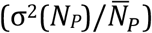, while the coefficient of variation is equal to the standard deviation divided by the mean 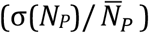. Both measures provide information about the dispersion of the distribution of expressed molecules relative to its mean.

The Fano factor and the coefficient of variation display the same behavior as the standard deviation in the number of expressed molecules (i.e., the size of the fluctuations). When a gene is highly induced at the highest affinity value, residence time does not affect any of these quantities (Sup Fig. 1). As explained in the main text, the reason is that the regulated gene behaves like a constitutive gene in this case, where the fluctuations of protein concentrations around their mean do not depend on a TF’s residence time (Fig. 2a and d; Raj and van Oudenaarden 2008).

At the opposite extreme, as induction approaches zero, longer residence times increase the Fano factor and the coefficient of variation. The reason is that longer residence times produce larger fluctuations in the number of expressed molecules (Fig. 2a). Not surprisingly then, the variance in protein number is higher for longer residence times, which increases the strength of noise (Sup Fig. 1), because the mean protein and mRNA expression is not affected by residence time (Fig. 2e; Sup Fig. 2e). In sum, similar to our results in the main text, as the level of induction decreases, the amount of noise decreases with residence time.

### 3. Protein and mRNA production and degradation events

Because shorter residence times (at submaximal induction) reduce variation in the frequency of protein production and degradation events, we hypothesized that production and degradation events should alternate more regularly at short residence times, such that the number of consecutive production events (i.e., the number of protein production events without any intervening degradation event should decrease). To validate this hypothesis, we quantified the number of consecutive production events at different levels of induction and residence times. When induction is high, residence time does not affect the number of consecutive production events (Sup. Fig. 3c and e), because gene expression resembles that of a constitutive gene at all residence times (Fig. 2a and b). However, as induction decreases, shorter residence times decrease both the mean and standard deviation number of consecutive production events (Sup Fig. 3c-f), showing that production and degradation events alternate more frequently.

One exception to this pattern occurs at the lowest level of induction. Here, the mean and standard deviation of the number of consecutive production events reaches a maximum at intermediate residence time and decreases as the residence time increases (Sup Fig. 3c-f, yellow). We believe that the reason is that most production events tend to occur in a few but long periods of active gene expression when the induction level is low and the residence time is long. During these long periods of active gene expression, protein expression behaves as for a highly induced gene, which has a smaller mean and standard deviation of the number of consecutive production events (Sup. Fig. 3c-f). Notice, however, that this dynamic will still produce high levels of noise, because mRNA and protein molecules will fluctuate between very low and very high levels (Fig 2a; Sup Fig. 2a). In conclusion, the more frequent alternation of production and degradation events observed at shorter residence times produce a more regular and less noisy dynamic of expression.

